# Driving the scalability of DNA-based information storage systems

**DOI:** 10.1101/591594

**Authors:** Kyle J. Tomek, Kevin Volkel, Alexander Simpson, Austin G. Hass, Elaine W. Indermaur, James Tuck, Albert J. Keung

## Abstract

The extreme density of DNA presents a compelling advantage over current storage media; however, in order to reach practical capacities, new approaches for organizing and accessing information are needed. Here we use chemical handles to selectively extract unique files from a complex database of DNA mimicking 5 TB of data and design and implement a nested file address system that increases the theoretical maximum capacity of DNA storage systems by five orders of magnitude. These advancements enable the development and future scaling of DNA-based data storage systems with reasonable modern capacities and file access capabilities.

## MAIN

DNA is an excellent candidate for archival data storage as it offers high raw information density as well as durability and energy efficiency^1–4^. Dense DNA databases will be highly diverse, crowded, and physically disordered, thus posing inherent challenges to data organization and retrieval. Existing systems have been 200 MB or less in capacity; these data volumes can be completely read by modern DNA sequencing technologies^3,5–12^. In contrast, high-capacity systems will not be able to be sequenced in their entirety (**Fig. 1, Supp. Fig. 1**), nor will entire databases be able to be decoded and stored using low latency systems that are higher in storage hierarchies. High-capacity systems will also require a large number of available file addresses (i.e. PCR primer sequences^5,7–12^), to organize the data. However, due to potential off-target molecular interactions, addresses must be sufficiently different from each other in sequence and are, therefore, finite in number and limit total system capacities (**Fig. 1, Supp. Fig. 1**).

**Figure 1.**
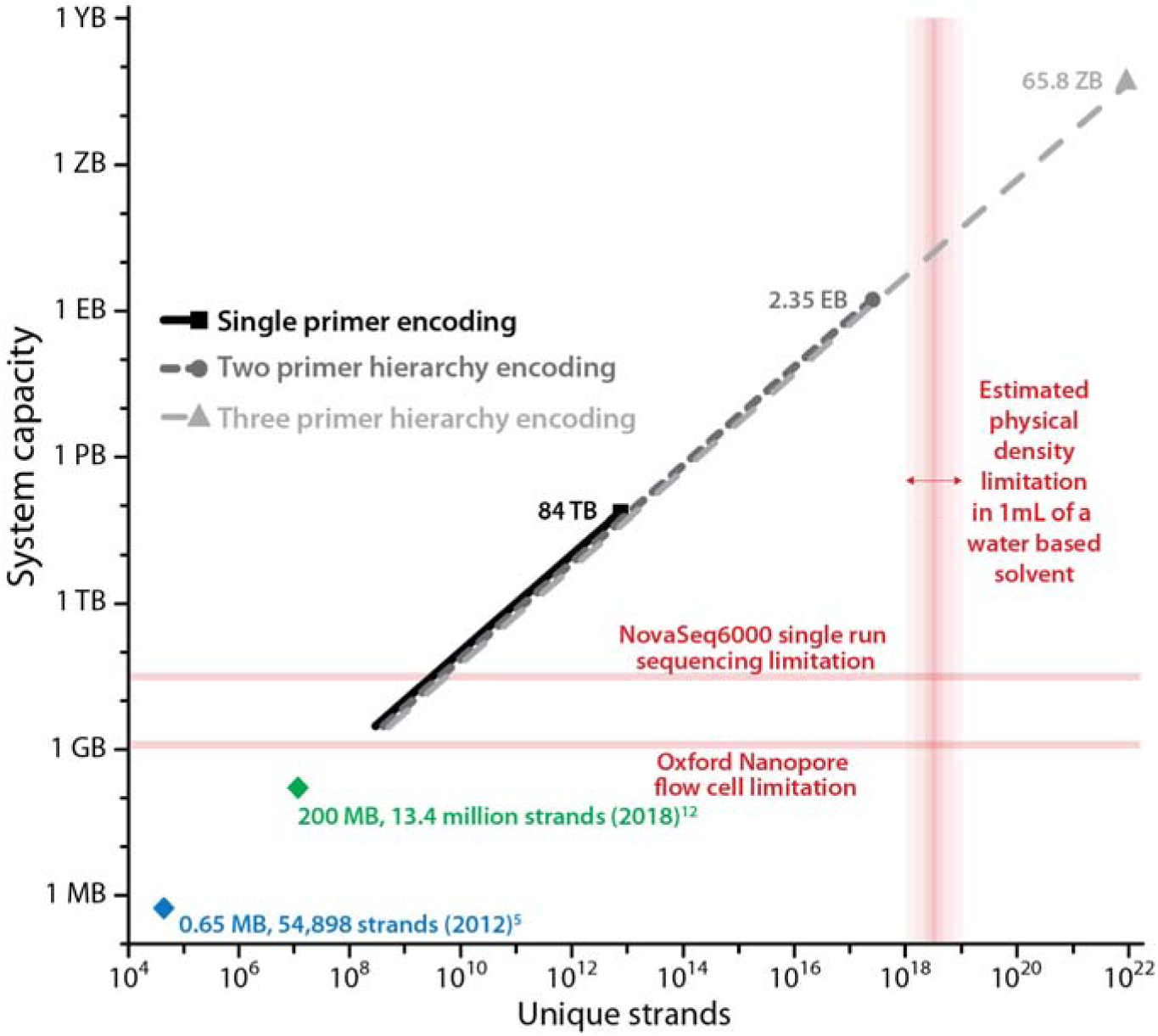
Theoretical analysis of readable files sizes, total system capacity limits, and improvements through physical data extraction and nested encoding. *Limited readable files sizes:* Current sequencing platforms can only sequence a fraction (~ 20-30 GB) of the theoretical maximum capacity of current systems (84 TB) assuming a sequencing depth of 10. The limits of commonly used next generation sequencing platforms are included for reference: Oxford nanopore flow cells can sequence 1.5×10^11^ bases, or roughly 1.27 GB per flow cell using our encoding scheme and average sequencing depth of 10. Illumine’s Novaseq6000 platform can sequence 2×10^10^ of our 200 bp strands per run, or roughly 28.9 GB. *Capacity limits*: The linear plots of system capacities are based on current best estimates of 28,000 usable primers ^12^ and an average file size stored per unique address of 3 GB. As the total number of unique strands within a database increases, so does the total system capacity, limited ultimately by the number of primers available. Thus, the availability of non-interacting primers limits the theoretical maximum capacity of storage systems. The system capacity limit for current one-primer encodings using 28,000 primers (all 27,999 files sharing 1 antisense primer) storing 3-GB files is 84 TB; this corresponds to 7.88×10^12^ unique 200 bp long strands. In contrast, using the same distinct primers in double or triple nested architectures increases the number of possible addresses exponentially. As a result, the total capacities also increase to 2.35 EB (2.52×10^17^ unique strands) and 65.8 ZB (8.98×10^21^ unique strands), respectively. The aqueous solubility of DNA is roughly between 10^18^ and 10^19^ per milliliter, depending on ionic concentrations.

Here we present a platform for accessing specific data from high-capacity DNA-based databases in conjunction with a nested file address system that can handle the organization of exascale databases. We will refer to this overall storage system, which uses DNA Enrichment and Nested SEparation, as DENSE data storage. This system directly addresses the challenges arising from the molecularly crowded nature of high-capacity DNA storage systems while functioning within a single physical pool of DNA. Therefore, it not only harnesses the raw capacity and density advantages of DNA but also drives the practical scalability of high-capacity data storage systems.

The current state-of-the-art file access method uses polymerase chain reaction (PCR) to amplify a desired file’s corresponding DNA strands (referred to as random access^7,8,11,12^). However, random access is theoretically predicted to exhibit decreasing sequencing efficiencies with increasing database size when PCR will not be able to overwhelm large quantities of non-target database strands. To experimentally measure this transition point, we generated a library of five files, each with unique PCR primer sequences (**Fig. 2a, Supp. Fig. 2a**), and mixed it with increasing quantities of background database strands. As it is currently cost prohibitive to order large databases of completely unique strands of DNA, large DNA databases can be mimicked in mass proportions by mixing copies of an individual file (i.e. 1.94 GB of File 3 strands = 1.14E9 total strands) with many more background database strands (i.e. 6.22 GB to 19.4 TB of a single non-specific DNA strand, 3.66E9 to 1.14E13 total strands, respectively). After 30 cycles of random-access PCR to amplify File 3 in this series of databases, the relative abundance of File 3 strands to background DNA was compared by quantitative PCR. As predicted, the percentage of the sample that was File 3 monotonically decreased as a function of increasing background DNA (**Fig. 2b**). File 3 fell below 50% of the total sample once the database size reached 31.1 GB and higher. Thus, in high-capacity systems random access becomes ineffective for specific file retrieval.

**Figure 2.**
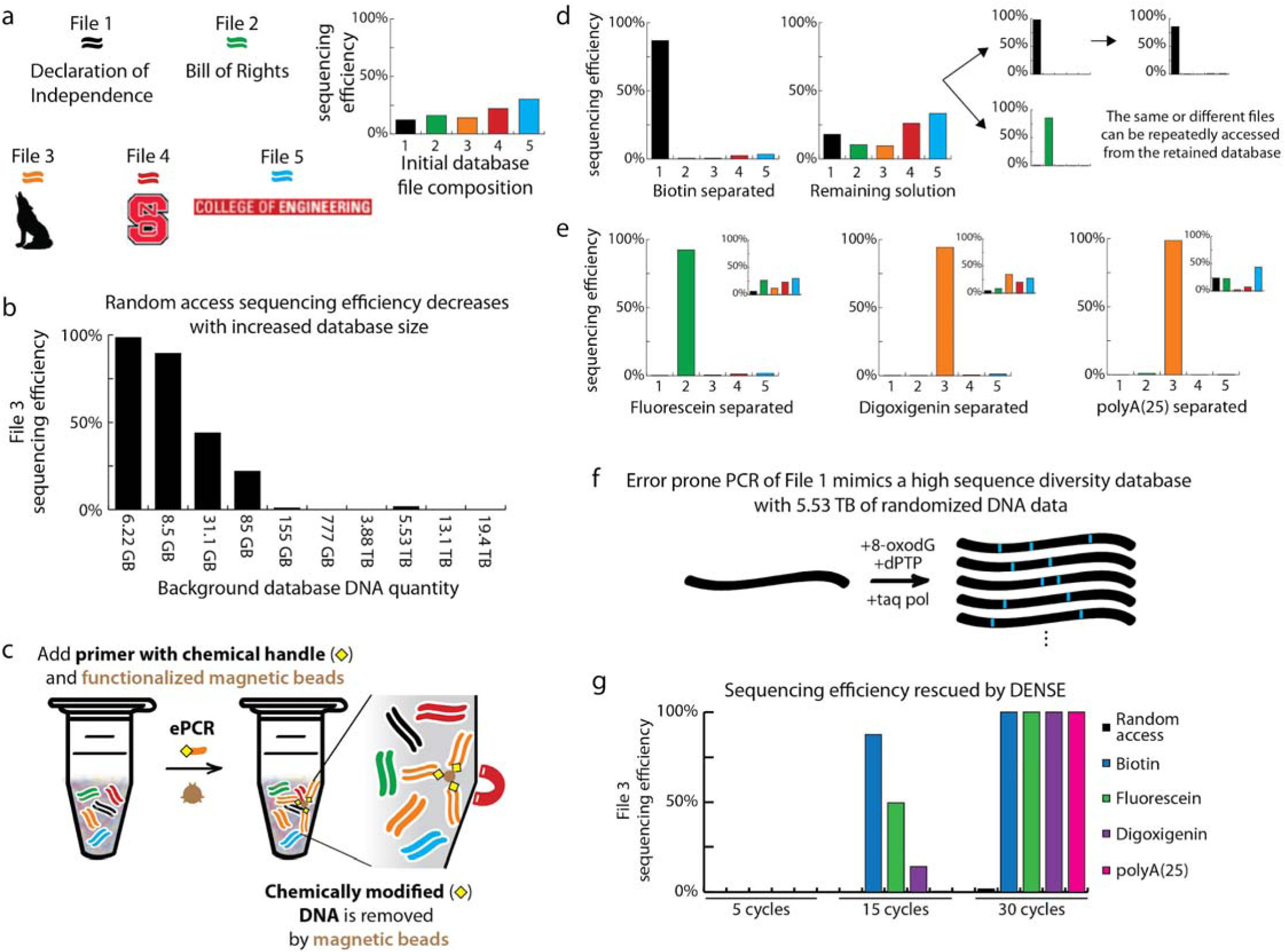
Physical file separations in DENSE storage rescue the decreased sequencing efficiency experienced by high-capacity databases. **a**, A library of five files was ordered and analyzed using NGS to confirm an even file distribution. **b**, File 3 strands were enriched over increasingly higher capacity backgrounds of non-specific DNA strands using 30 cycles of random-access PCR. Random access failed to enrich File 3 to above 50% of the total sample once the background capacity reached 31.1 GB, as measured by quantitative PCR. **c**, DENSE physically extracts a file (orange) from the database so only its strands are sequenced. A primer functionalized with a chemical handle (yellow diamond) is used to execute one emulsion PCR cycle to create chemically labeled copies of the desired file’s strands. Functionalized magnetic beads (brown) that bind to the chemical handle are added to the sample. The desired file is bound to the bead, and the unbound solution containing the original database is removed and saved for future reuse. The bound file is then eluted from the bead. **d**, After biotin-streptavidin file extractions, the remaining solution still contained all files while the target files were enriched and physically separated, as measured by next generation sequencing. By mapping sequencing reads to the original file sequences, all targeted data were confirmed recovered. The target file was retained in the supernatant containing the database and was able to be copied and extracted again. File 1 was extracted three sequential times, and File 2 was extracted from the solution remaining after an initial extraction of File 1. **e**, File extractions using fluorescein, digoxigenin, and polyA(25) as chemical handles also successfully separated target files from the database. **f**, A large-scale background mimicking diverse data was created using error prone PCR ^13^ to mutagenize and amplify File 1. **g**, Random access was compared directly to chemical handle extractions. File 3 strands, with a starting fraction of 0.03% of the total number of strands, were enriched over a high-capacity background equivalent to 5.53 TB of undesired, non-specific strands using either random access (black) or PCR followed by chemical handle primer extractions (blue, green, purple or pink). After 5, 15, and 30 cycles of PCR (random access), enrichment of File 3 was 0.0%, 0.0%, and 1.69% of the total sample, respectively. After biotin-modified PCR followed by extraction, the enrichment of File 3 was 0.2%, 87.5%, and 100% of the total sample, respectively. After fluorescein-modified PCR followed by extraction, the enrichment of File 3 was 0.1%, 49.6%, and 100% of the total sample, respectively. After digoxigenin-modified PCR followed by extraction, the enrichment of File 3 was 0.2%, 14.2%, and 100% of the total sample, respectively. After poly(A)-25-modified PCR followed by extraction, the enrichment of File 3 was 0.09%, 0.47%, and 100% of the total sample, respectively.

To address this database capacity limitation, we sought to physically separate specific files from the database, allowing for the efficient sequencing and analysis of only desired data. To this end, PCR primers modified by chemical moieties were used to create chemically labeled copies of a desired file’s DNA strands (**Fig. 2c**). These copies of individual files were then separated from the database of five files using magnetic beads and fully recovered as confirmed by sequencing (**Fig. 2d, e, Supp. Fig. 2b-d**). Four distinct modification systems were capable of efficient and complete file access (biotin-streptavidin, fluorescein-antibody, digoxigenin-antibody, polyA-polyT oligomers). Next generation sequencing results indicate a sequencing efficiency above 86%, representing a reduction in wasted sequencing throughput. Of note, to access files in this manner, we found that only a single emulsion PCR cycle was needed to chemically label files. Importantly, we observed no destruction of the original database in the remaining solution following separation (**Fig. 2d, e, Supp. Fig. 2d**). Furthermore, we determined that the same or a different file could be repeatedly accessed from this previously ‘used’ solution (**Fig. 2d**). Taken together, our approach to physically separate files is non-destructive and represents a reusable DNA-based storage system.

To directly compare the performance of DENSE storage with random access in high-capacity systems, we compared the relative enrichments of File 3 from a 5.53-TB database (**Figure 2f, g**). In this experiment, to better mimic a true high-capacity and high-diversity database, File 1 was mutagenized by two rounds of error prone PCR^13^ to an estimated 5.53 TB of unique data. Whereas random access was not able to significantly enrich File 3 strands from this high-capacity database, all four DENSE separation methods enriched File 3 to above 99% of the total sample after 30 cycles of modified emulsion PCR.

High-capacity systems require many unique addresses in order to store and access information, yet there are fewer than 30,000 usable primer addresses that will not cross interact^12^. For instance, in a storage system comprised of 3 GB file sizes, 30,000 primers limit total system capacity to ~84 TB (**Fig. 1, ‘Single primer encoding’**), given our strand organization and encoding strategy. To address this database capacity limitation, DENSE uses a hierarchical encoding scheme where primer sequences are nested and used in sequential combination (**Fig. 3a**)^14^. Theoretically, this architecture can more than exponentially increase the number of unique addresses for files without increasing the total number of unique primers needed: nesting 2 primers would enable exascale capacities (**Fig. 1, ‘Two primer hierarchy encoding’**), while nesting more than 2 primers would result in exponentially larger numbers of total addresses (i.e. 30,000 unique primers^N number of nests^). Using this hierarchical PCR architecture with nested primers used in sequential combination (but with no physical extractions), both File 4 and File 5 were separately and selectively accessed using opposite temporal amplification sequences, albeit there were substantial amounts of contaminating off-target strands (**Fig. 3b, Supp. Fig. 3**). Combining this hierarchical strategy with biotin separations resulted in a reduction of these contaminating off-target strands. Specifically, the desired file in each case comprised either 96.9% or 86.8% of the sample, as measured by quantitative PCR, showing the specificity of nested addresses when used in the correct hierarchical temporal sequence and in conjunction with file separation (**Fig. 3b, Supp. Fig. 3**).

**Figure 3.**
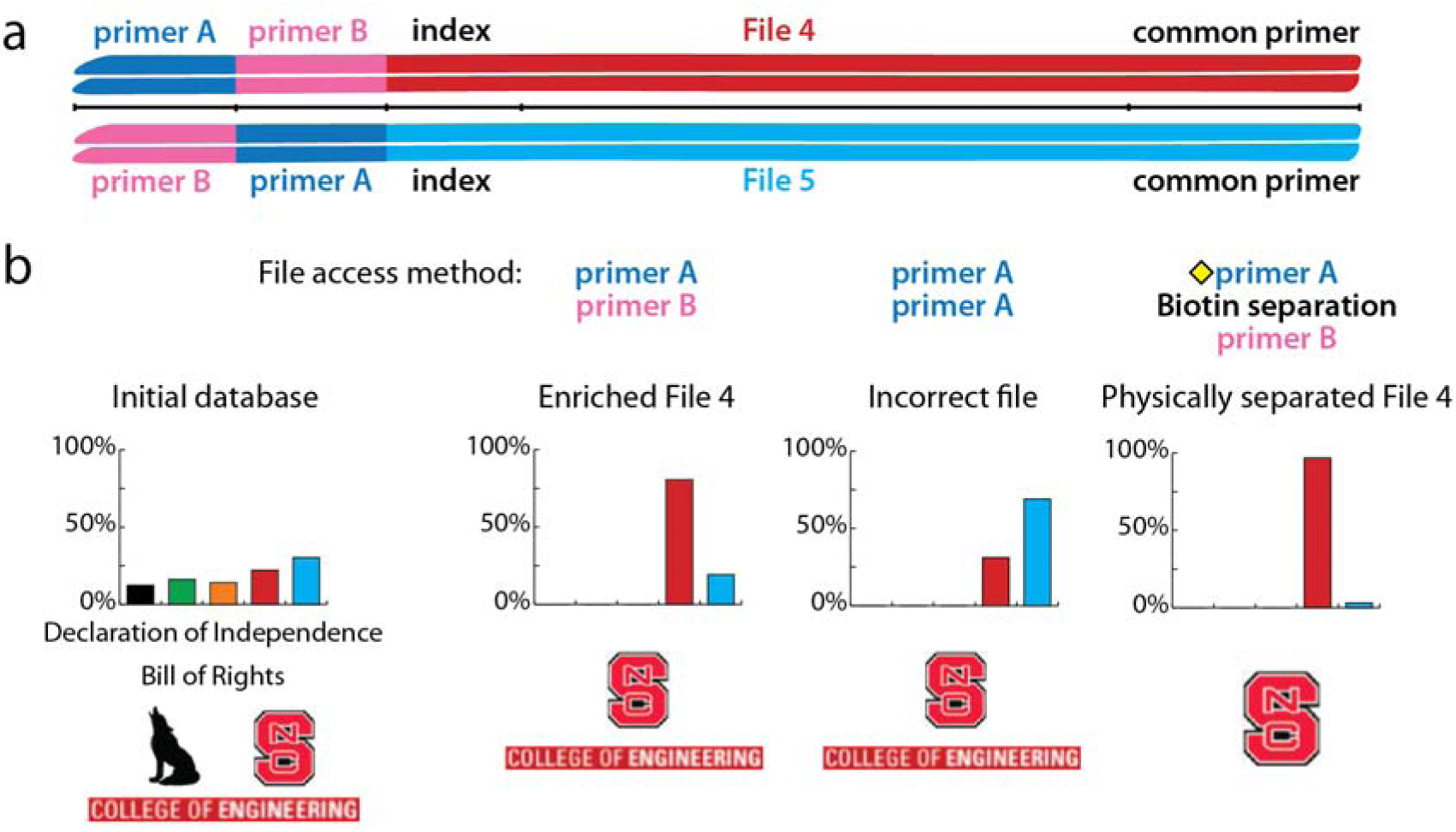
Combining a nested, hierarchical address strategy with physical separations results in purified enrichment of the desired file. **a**, Strand architectures of Files 4 and 5 exhibit nested primer addresses. Binding sites for primers A and B are shared by both files but in opposite orders. Both files share a common antisense primer. **b**, Experimental demonstration that PCR using primer A followed by primer B enriches for File 4. PCR amplifications using two rounds of the same primer enriches for the incorrect file. In conjunction with physical extractions, File 4 is specifically accessed using hierarchical PCRs. The extraction after the first PCR amplification increases File 4 enrichment from 81% to 97% over no extraction, as measured by qPCR.

DENSE is practical in that it reduces the number of PCR cycles needed compared to random-access methods: only 1 cycle was necessary to access data from the 5-file database. This not only reduces the amount of dNTPs and other reagents needed, but also reduces the chances of mutational errors and alterations in strand distributions that may arise from PCR (**Supp. Figs. 4-6**). Consequently, encoding algorithms may not need to sacrifice as much information density towards error correction. We do note that when the capacity of databases increase, more PCR cycles are required to access target files using DENSE (**Fig. 2g**). At higher capacities DENSE outperforms PCR alone (random access was not able to access target files at all), but this requirement for increased PCR cycles suggests additional biochemical engineering should be pursued to improve the specificities and affinities of the many complex molecular interactions that can occur during file separation.

Although initially designed to address barriers to scaling to extreme capacities (~PB and higher), DENSE storage will be useful at modern GB or TB capacities that are routinely achieved using mainstream DNA synthesis technologies. For example, if common file sizes of ~25 MB are desired, there will be challenges in providing enough unique addresses for GB to TB level systems without harnessing DENSE storage’s nested address architecture (**Supp. Fig. 1, ‘25 MB files’**). Furthermore, without physical file separations, reading data from GB to TB level systems will be wasteful and perhaps infeasible even using state-of-the-art sequencing capabilities. For instance, Illumina’s NovaSeq6000 can read only 20-30 GB of data when conservatively accounting for 10 redundant copies per strand (i.e. read depth of 10) (**Fig. 1, Supp. Fig. 1**). Critically, this work demonstrates the enrichment and physical separation of the equivalent of 1.94 GB of targeted DNA strands from 5.53 TB of undesired database strands (**Fig. 2g**). This approach can be combined with a hierarchical, nested-address system to increase the theoretical total capacity of DNA storage systems by over five orders of magnitude.

While there are many challenges, and likely many still unanticipated, there are recent promising breakthroughs in all necessary aspects of DNA storage: advances continue to be made in DNA synthesis and sequencing, in encoding and error correction, and now in physical file access and system architecture. This work provides a conceptual and quantitative framework to think about DNA storage systems and their challenges, proposes practical strategies to address key barriers to scaling system capacities, and suggests that DNA-based data storage systems with reasonable modern capacities and file access capabilities are not only immediately achievable but also scalable to extreme capacities in the future.

## METHODS

### Data Representation, Encoding, and Decoding

We adopted a similar approach for representing and encoding data as reported in recent work^6,8,12^. We partitioned a digital file into blocks of data that fit in DNA strands that are 200 bp long. Each strand consists of multiple fields. A primer binding site occupies each end and enables DNA polymerase chain reactions. Between the primers, we placed three fields that represent the index of the strand within the file, the data payload, and a checksum to detect errors within the strand. We used a fixed length index that is 16 bp-long. This leaves the remaining 144 bp-long sequence to represent the data payload of each strand. We designed 8 bp-long codewords to represent one byte of data. The codewords have no repetition of bases both individually and when appended, and they are GC balanced. Each byte of file is converted one byte at a time into a corresponding codeword and appended together to form the payload of a strand. The checksum is a single-byte XOR-accumulation of all the data in the payload that is encoded and appended to the end of the data payload. The checksum allows each strand to self-check its own data. The only notable difference for hierarchical encoding is that it requires an additional primer in each strand, thereby reducing the size of the data payload.

We also adopted a redundant XOR-style encoding proposed by Bornholt et al.^8^ to enhance the reliability of our system. In our design, indices with even values hold data, and odd indices store the XOR-ed content of their adjacent strands. This redundancy enables recovery of data even if some strands are lost or discarded due to an invalid checksum. The decoder algorithm for our encoding is similar to previous work^8^, with the modification that we can disregard any read with an invalid checksum. It is important to note that for clarity of analysis and ease of comparison across systems, the file and database sizes estimated in the figures do not take into account the overhead required to implement XOR or other encodings that may be used. Thus, we present best case scenarios, whereas true capacity challenges and limitations are likely even more severe than described in this work.

### Primer design

Primers used in this work were designed to achieve multiple goals. First, they must facilitate effective PCRs. The primers were designed such that GC content is between 40% and 60%, and their melting temperature is between 50°C and 60°C. We required that the last base is G but the GC content in the last 5 bases could not exceed 60%. Second, primers were designed to reduce the likelihood of non-specific binding with other primer binding sites. We required a Hamming distance of >10 between all primers to minimize the likelihood of such binding. We also performed NUPACK simulations of homodimer, hairpin, and heterodimer bindings^15^. We required a Gibbs free energy greater than −10 kcal/mol at 50°C on all likely complexes to select the primer. Note, we compared each candidate primer to all other primers to ensure no heterodimer bindings are likely, and we included the Illumina NEXTERA primers in this process. Third, to reduce the likelihood of non-specific binding between a primer and the data payload, we required that primers must contain a repeating nt every 5 bases. This guaranteed that primers would differ from all length 20 sub-sequences of the data payload.

We used a computer program written in Python to automate the generation of candidate primer sequences and screened them against the requirements stated above. The python program invoked the relevant analysis in NUPACK as needed.

### Emulsion PCR

The emulsion PCR (ePCR) protocol from Schutze et al.^16^ was modified slightly and used for all PCR steps. Emulsions were created by mixing 150 μL of emulsion oils (73% Tegosoft DEC (Evonik, 99068594), 20% mineral oils (Sigma Aldrich, 330779), 7% ABIL WE (Evonik, 99068358)) with 25 μL aqueous PCR samples. Samples were then vortexed for 5 minutes until a persistent emulsion was formed. Samples were aliquoted into four PCR tubes and a standard Q5 polymerase PCR protocol was used. Twenty cycles were sufficient to reach the maximum yield of DNA product. After amplification, aliquots were pooled in an Eppendorf tube and emulsions were broken with the addition of 1 mL of isobutanol followed by a 5 second vortex. Five volumes of (125 μL for 25 μL PCR reaction volume) binding buffer (Biobasic Canada Inc. BS664) was added to samples, gently mixed and centrifuged at 2,400 g for 30 seconds. The organic phase was removed and discarded while the remaining aqueous phase was purified using AMPure XP beads (Beckman Coulter, A63881). DNA was eluted in 50 μL of water.

### Biotin-Streptavidin file extractions

File-specific sense (‘coding’) primers were ordered with a biotin modification on the 5’ end. PCR amplified samples were purified (AMPure XP beads) and added to prewashed streptavidin magnetic beads (NEB #S1420S) (wash and bind buffer: 20 mM Tris-HCl pH 7.4, 2M NaCl, 2 mM EDTA pH 8) and incubated at room temperature on a rotisserie for 30 minutes. The database files were retained by collecting the supernatant. The beads were then washed once with 100 μL of the binding buffer and once with 100 μL of a low-salt wash buffer (20 mM Tris-HCl pH 7.4, 150mM NaCl, 2 mM EDTA pH 8). Amplified DNA was subsequently eluted (elution buffer: 95% formamide (Sigma, F9037) in water). DNA sizes and concentrations of the purified (AMPure XP beads) supernatants and elutions were measured on a Fragment Analyzer (Advanced Analytical, DNF-474) before the addition of Illumina sequencing adapters. Representative DNA gel images of biotin separations are shown in **Supplementary Figure 2b.**

### Fluorescein and digoxigenin file extractions

File-specific sense (‘coding’) primers were ordered with either fluorescein or digoxigenin on the 5’ end (Eurofins Genomics). Antibodies (anti-fluorescein: Novus Biologicals, NB600-493, Lot 19458; anti-Digoxigenin (21H8): Novus Biologicals, NBP2-31191, 17E16) were bound to magnetic protein A or G beads (BioRad Cat. #s 161-4013 & 161-4023) through a 30-minute room temperature incubation (bind and wash buffer: 20 mM Tris-HCl pH 8, 300 mM NaCl, 2 mM EDTA). PCR amplified samples were purified (AMPure XP beads) and added to the antibody-linked beads and incubated at room temperature on a rotisserie for 2 hours. The database files were retained by collecting the supernatant. The beads were washed once with 100 μL of the binding buffer and once with 100 μL of a low salt wash buffer (20 mM Tris-HCl pH 7.4, 150mM NaCl, 2 mM EDTA pH 8). DNA sizes and concentrations of the purified (AMPure XP beads) supernatants and elutions were measured on a Fragment Analyzer (Advanced Analytical, DNF-474) before the addition of Illumina sequencing adapters. Representative DNA gel images of a fluorescein separation are shown in **Supplementary Figure 2c.**

### Oligo-d(T) magnetic bead separation

File-specific sense (‘coding’) primers were ordered with a poly(A)-25 tail on the 5’ end (Eurofins Genomics). Poly-d(T) beads (NEB #S1419S) were washed twice with 100 μL wash and bind buffer (20 mM Tris-HCl pH 7.4, 2M NaCl, 2 mM EDTA pH 8). PCR amplified samples were purified (AMPure XP beads) and added to the desired amount of bead based on the amount of DNA present and theoretical binding capacity. The mixture was heated in a thermal mixer at 90°C and 500 rpm for 2 minutes, allowed to cool to room temperature, and the database files were retained by removing the supernatant. The beads were washed twice with 100 μL of a low salt wash buffer (20 mM Tris-HCl pH 7.4, 150mM NaCl, 2 mM EDTA pH 8). Beads were then resuspended in 1x TE buffer, heated in the thermal mixer at 50°C and 500 rpm for 2 minutes. The desired file was extracted while the mixture was still hot by removing the eluted sample from the beads. DNA sizes and concentrations of the purified (AMPure XP beads) supernatants and elutions were measured on a Fragment Analyzer (Advanced Analytical, DNF-474) before the addition of Illumina sequencing adapters.

### Calculation of data quantity from total number of DNA strands

In Figures 1, 2, and Supp. Fig. 2 we refer to file and database sizes (MB, GB, etc.). For clarity and ease of comparison all values were calculated based on the total number of DNA strands. Each strand is comprised of 200 nts, 20 of which are used for each primer sequence, 16 for the index, and 8 for the checksum. 8 nts comprise each 1-byte codeword. Thus, each strand addressed with a single primer pair contains 17 bytes of data. Specifically, in Figure 2, we assumed a 10-copy physical redundancy per unique strand to provide a conservative estimate for a realistic system where multiple copies of each strand would likely be needed to avoid strand losses and inhomogeneous strand distributions. Thus, in Figure 2 total file and database sizes are divided by 10.

### Error prone PCR

Template DNA was amplified using 0.5 μL of Taq DNA polymerase (5 units/μL, Invitrogen, 100021276) in a 50 μL reaction containing 1X Taq polymerase Rxn Buffer (Invitrogen, Y02028), 2 mM MgCl2 (Invitrogen, Y02016), the sense and antisense primers at 1E13 strands each, and dATP (NEB, N0440S), dCTP (NEB, N0441S), dGTP (NEB, N0442S), dTTP (NEB, N0443S), dPTP (TriLink, N-2037), 8-oxo-dGT (TriLink, N-2034), each at 400 mM. PCR conditions were 95°C for 30 seconds, 50°C for 30 seconds and 72°C for 30 seconds for 35 cycles with a final 72°C extension step for 30 seconds.

### qPCR

qPCR was performed using SsoAdvanced Universal SYBR Green Supermix (BioRad). qPCRs were performed in 5 μL format using SYBR Green (95°C for 2 min, and then 50 cycles of: 95°C for 10 s, 50°C for 20 s, and 60°C for 20 s). qPCR results were compared to next generation sequencing results for samples that were analyzed using both methods. File compositions measured using both methods showed strong agreement (**Supp. Table 1**).

### Illumina library preparation

Illumina TruSeq Nano DNA Library Preps (Illumina, 20015965) were performed according to manufacturer instructions beginning from the ‘Repair Ends and Select Library Size’ step, as DNA fragmentation was unnecessary. The quality and band sizes of libraries were assessed using the High Sensitivity NGS Fragment Analysis Kit (Advanced Analytical, DNF-474) on the 12 capillary Fragment Analyzer (Advanced Analytical) at multiple steps during each protocol, typically after size selection and after PCR amplification. Unless otherwise stated, libraries were normalized to balance estimated sequencing depth across similar samples (e.g. all elutions had estimated sequencing depth of ~100 reads) using the molar concentrations measured on the Fragment Analyzer. The pooled sample had a concentration of 8 nM and was sequenced using the MiSeq v2 chemistry 150 PE kit that was operated as a 300 SR run. PhiX DNA was added at 20% of total DNA to increase sequence diversity.

### Error Analysis

Before proceeding with an error analysis of sequenced strands, the error-free reference strand for each sequenced strand needed to be determined. To find the error-free reference strands, a mapping operation was performed to match each sequenced strand with its original database strand. Due to the large number of sequenced strands in samples (up to 571k reads), the mapping operation was carried out in two steps: the first step partitioned the large read space using the primer sequences of the different files, and the second step further analyzed each partition to match each strand in a partition with its corresponding database strand.

The first step of mapping divided the initial sequencing read space into partitions, one for each file in the database, with the exception of Files 4 and 5 (hierarchical encodings) where each of these files had 2 partitions. These 2 partitions were used to separate nested address strands that were truncated from the first PCR step and reads where the nested address strands were not truncated. Other partitions were also created for special strands like the background strands used to simulate high-capacity data storage and for unknown strands that could not be categorized into a file’s partition. A strand from sequencing was placed into a partition by looking for a subsequence that matched a file’s sense primer, or the reverse complement of the anti-sense primer. The reverse complement of the anti-sense primer was used because all NGS sequencing reads are in the 5’ to 3’ direction. A subsequence was deemed acceptable if it matched a sense primer or anti-sense primer’s reverse complement within a Levenshtein distance of 4. A Levenshtein distance of 4 was chosen as the cut-off point to ensure that the matched subsequence was not data within a DNA strand, but one of the primers of interest. When a primer of interest is found in a sequenced strand, the sequenced strand is placed in the primer’s respective partition.

After categorizing each strand in a sample’s sequence pool, each partition was analyzed further to determine the original database strand for each sequenced strand in the partition. To find out the correct original strand, each original strand from a file was compared to each sequenced strand placed in the file’s partition by calculating the Levenshtein distance between the sequenced strand and the original strand. If the distance was less than or equal to 12, the original strand was considered as a candidate for a match. Because some of the original strands in the database have small edit distances between them, file strands that are close to the candidate were also checked against the sequenced strand to make sure the correct original strand was chosen. Once a candidate was concluded to correspond to a specific original strand, the location of the matching strand in the file along with the sequenced strand’s location in the read space was recorded. A distance of 12 was chosen as the threshold in order to reduce the amount of checking that was required once a candidate was found, while ensuring that error rates would not be artificially low due to choosing candidates that were within a small number of edit operations.

With a completed mapping of sequenced strands to their corresponding database strands, analyses such as error rates per base, strand error rates, and read distributions were performed. To calculate the error rate for a nt position, Equation 1 was used. Where L is the number of unique edit operations considered (insertions, deletions, substitutions), M is the number of unique strands in the database, s_j_ is the jth strand in the database, N_j_ is the number of sequenced strands that map to strand s_j_, s_k_ is the kth strand that maps to database strand s_j_, T is the total number of strands from the sample that has been mapped to some database strand, and EO_l_(s_j_, s_k_)_i_ is the number of edit operations of type l at the ith nt position to transform s_j_ to s_k_. This equation calculates the total error rate for base position i by summing up all of the edit operations of each type at the ith position needed to transform each original database strand to the sequenced strands that map to it, and then dividing by the total number of mapped strands in the sample.

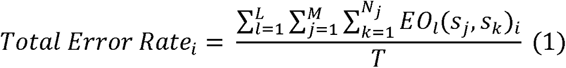

Similarly, the error rate for each strand in the original database was calculated using Equation 2. Where L is the number of unique edit operations, s_j_ is a strand from the original database, N_j_ is the number of sequenced strands that map to strand s_j_, s_k_ is the kth strand that maps to s_j_, T_sj_ is the total number of mappings in the sample for s_j_, and EO_l_(s_j_, s_k_) is the number of edit operations of type l to transform s_j_ to s_k_.

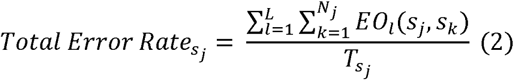

## ACKNOWLEDGEMENTS

We acknowledge Kevin N. Lin for helpful discussions. This work was supported by a National Science Foundation Grant #CNS-1650148 and a North Carolina State University Research and Innovation Seed Funding Award #2018-2509 to AJK and JT. KJT was partially supported by a Department of Education Graduate Assistance in Areas of Need Fellowship. EWI and AH were partially supported by funds from the North Carolina State University Research Experience for Undergraduates, Provost’s Professional Experience Programs, and NCSU startup funds.

## AUTHOR CONTRIBUTIONS

KJT, JT, and AJK conceived the study. KJT, EWI, and AJK developed the wet experimental system. KV, AS, and JT developed the software and simulations. KJT, EWI, AGH, and AJK planned and performed the wetlab experiments with guidance from all. KV, AS, and JT planned and performed simulations and next generation sequencing analysis with guidance from all. KJT, AJK, KV, and JT wrote the paper with input from all.

