## Supplementary data for "Driving the scalability of DNA-based information storage systems"

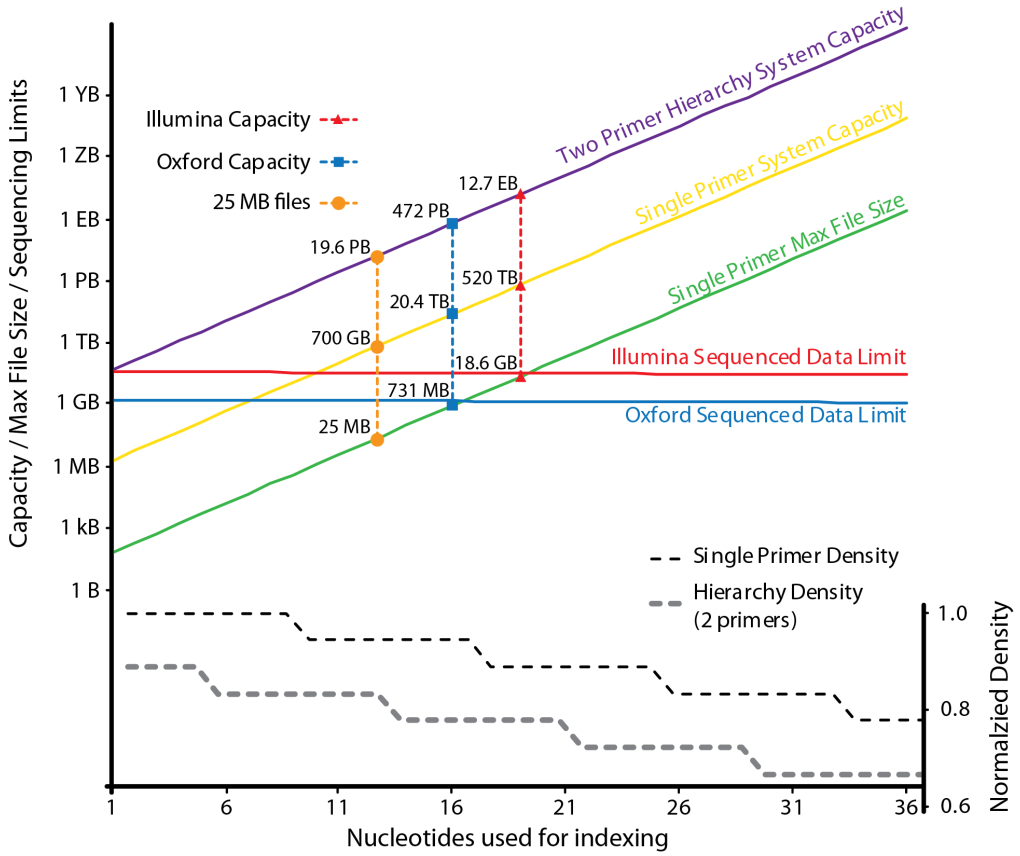


**Supplementary Figure 1. Implementation of high capacity DNA-based data storage presents physical and architectural challenges.** This plot presents a quantitative analysis of tradeoffs in selecting system parameters. The maximum file size as well as total storage capacities for both hierarchy (nested addresses) (purple line) and single primer (yellow line) systems increase with increasing number of nucleotides (nt) allocated to the index region of a strand (x-axis). Also plotted are the densities of single primer (black dash) and hierarchical (grey dash) strand architectures (normalized to the density of a single primer with 1 nt for indexing), the amount of data that can be sequenced with two different sequencing methods (Illumina – solid red line and Oxford Nanopore – solid blue line), and the maximum file size that can be attained as a function of index length (solid green line). The maximum amount of data that can be sequenced assumes a 10x sequencing depth. The maximum file size is plotted for only the single primer configuration. For clarity, the amount of data that can be sequenced and the maximum file sizes for hierarchy systems are not plotted but can be calculated using the provided densities and would be only minimally offset from the single primer system.

a

| File # | Strands | Size including encoding | Raw size excluding encoding overhead | Format | Encoded File |
|---|---|---|---|---|---|
| 1 | 876 | 2.34 kB | 14.89 kB | .txt | Declaration of Independence [Adopted in Congress 4 July 1776] The Unanimous Declaration... |
| 2 | 999 | 7.99 kB | 16.98kB | .zip | We the People of the United States, in Order to form a more perfect Union, establish Justice... |
| 3 | 1143 | 9.15 kB | 19.43kB | .png | |
| 4 | 1577 | 11.03 kB | 22.87 kB | .png | |
| 5 | 1389 | 9.72 kB | 20.14 kB | .png | COLLEGE OF ENGINEERING |
| | **5984** | **40.23 kB** | **94.31 kB** | | **TOTALS** |

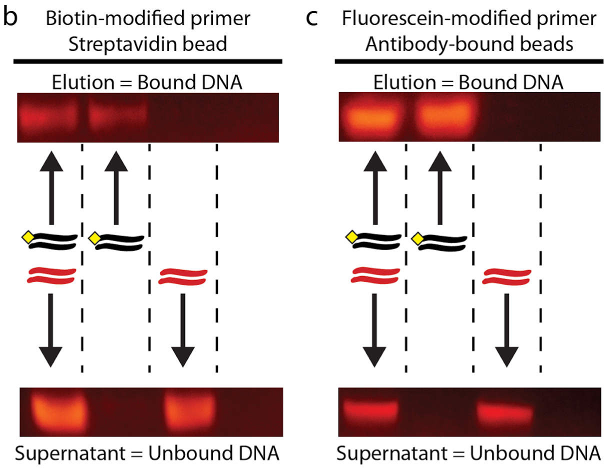


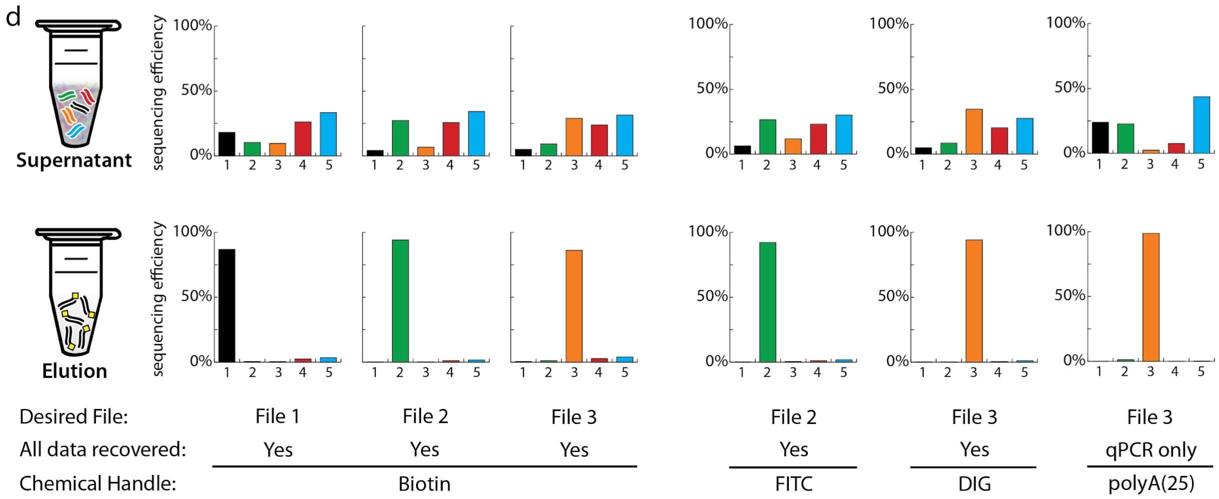


**Supplementary Figure 2. Library description, preliminary data, and complete analysis of file separations. (a)** Description of the experimental database. Five files totaling 40.23 KB (5,984 unique strands), were encoded, synthesized, pooled and stored as one database. **(b,c)** Proof of concept DNA strand extraction. Biotin **(b)** or fluorescein **(c)** primers were used to amplify a single 200 bp-long strand of DNA. Extraction reactions were performed with either: a mixture of modified (black with yellow diamond) and unmodified (red) DNA (lane 1), modified DNA (lane 2), unmodified DNA (lane 3) or water (lane 4). The resulting elutions and supernatants were visualized on an agarose gel. **(d)** After biotin-streptavidin, fluorescein-antibody, digoxigenin-antibody, and polyA(25)-Oligo-dT file extractions, the supernatants still contain all files (top) while the eluents were enriched for the target files (bottom), as measured by next generation sequencing. By mapping sequencing reads to the original file sequences, all targeted data were confirmed recovered.

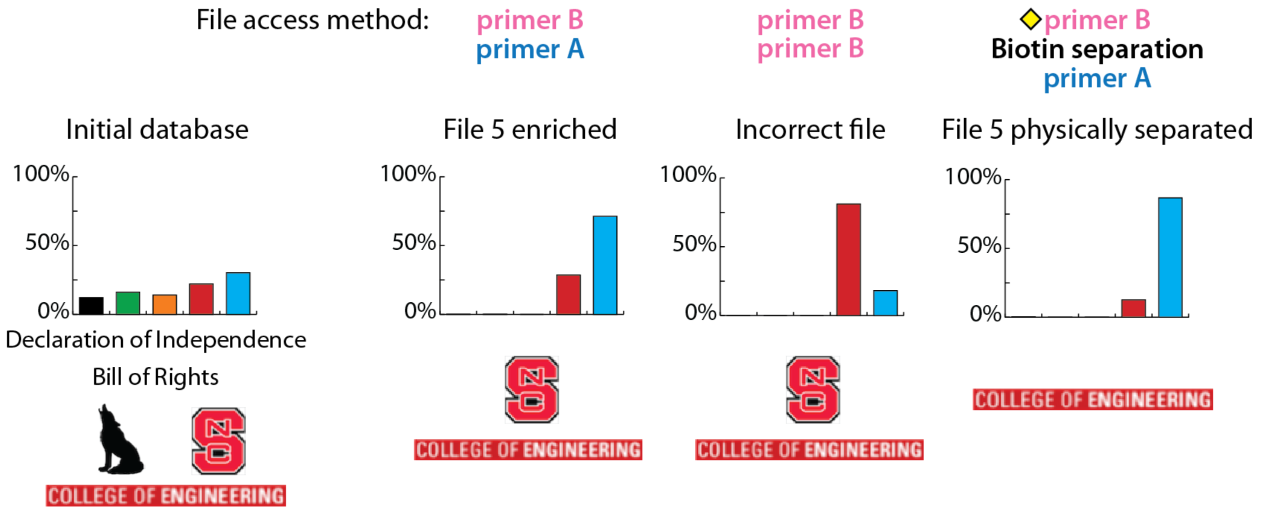


**Supplementary Figure 3. Combining a nested, hierarchical address strategy with physical separations resulted in purified enrichment of the desired file.** Experimental demonstration that PCR using primer B followed by primer A enriched for File 5. PCR amplifications using two rounds of primer B enriched for the incorrect file. In conjunction with physical extractions, File 5 was specifically accessed using hierarchical PCRs. The extraction after the first PCR amplification increased File 5 enrichment from 71% to 86% over no extraction, as measured by qPCR.

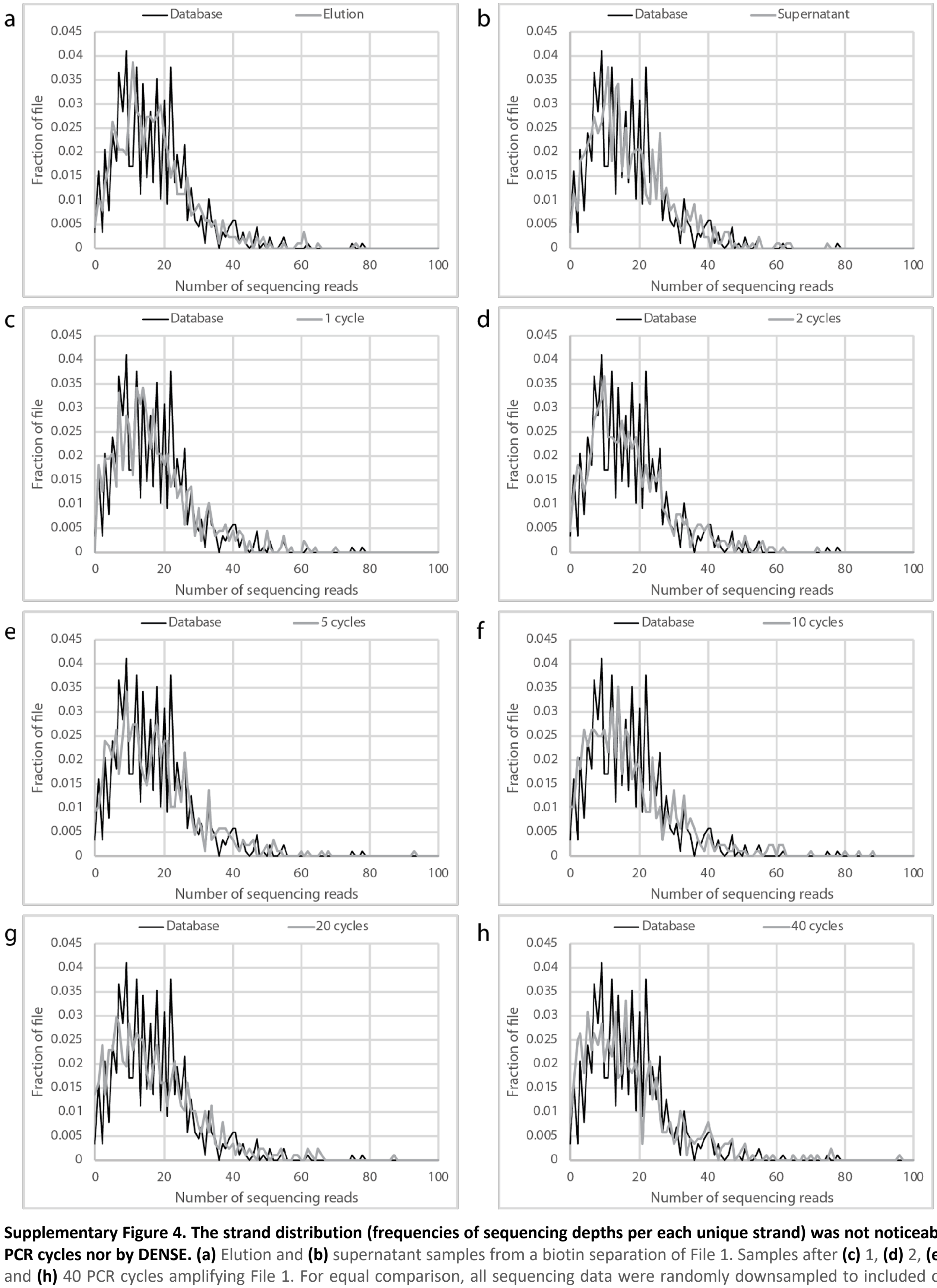


**Supplementary Figure 4. The strand distribution (frequencies of sequencing depths per each unique strand) was not noticeably affected by 40 PCR cycles nor by DENSE. (a)** Elution and **(b)** supernatant samples from a biotin separation of File 1. Samples after **(c)** 1, **(d)** 2, **(e)** 5, **(f)** 10, **(g)** 20, and **(h)** 40 PCR cycles amplifying File 1. For equal comparison, all sequencing data were randomly downsampled to included only 10,000 File 1 reads.

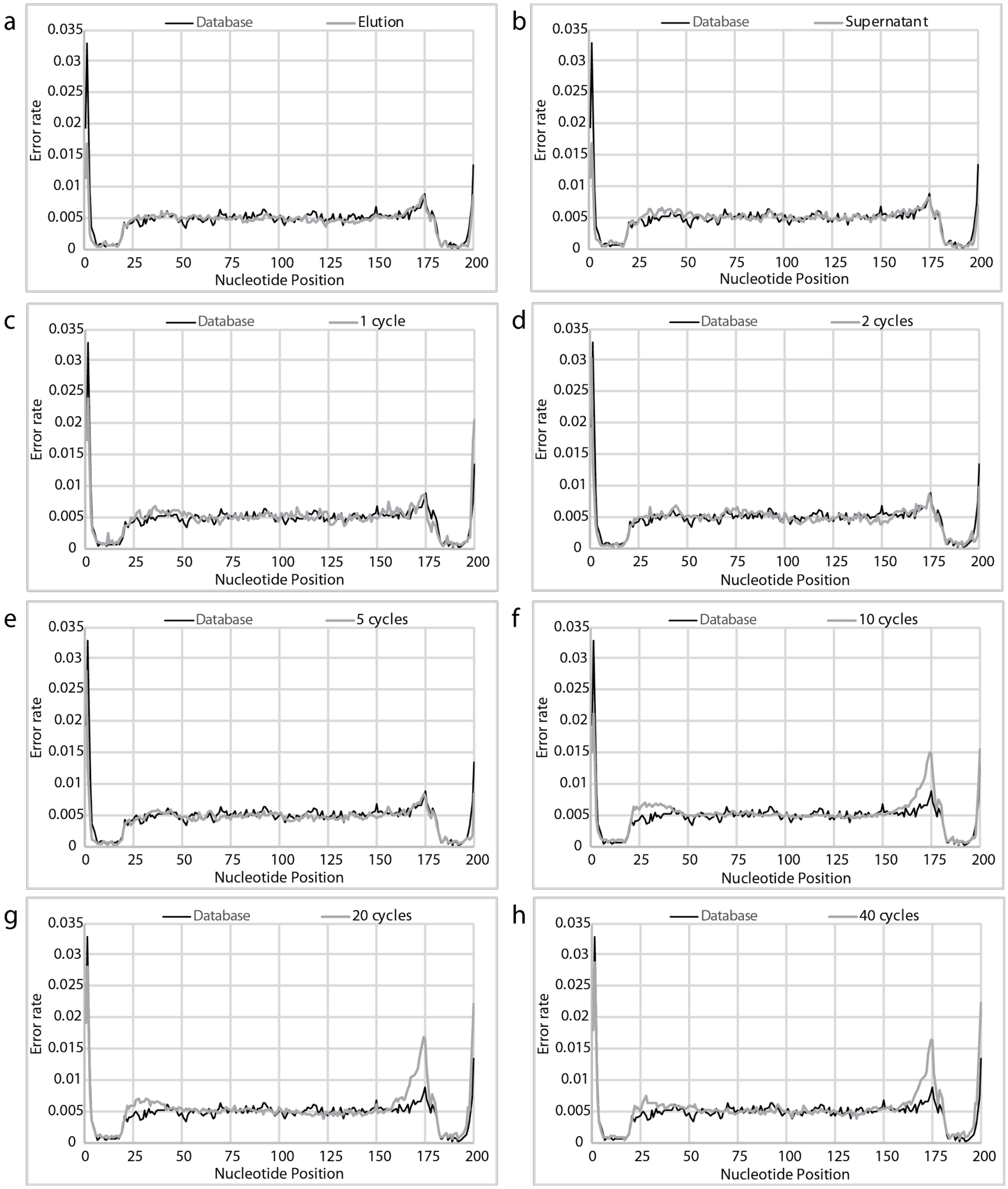


**Supplementary Figure 5. The error rate at a given nt position (1-200) is plotted as an average across all File 1 strands.** File 1 biotin separation **(a)** elution and **(b)** supernatant. **(c)** 1, **(d)** 2, **(e)** 5, **(f)** 10, **(g)** 20, and **(h)** 40 PCR cycles amplifying File 1. For equal comparison, all sequencing data were randomly downsampled to included only a random sample of 10,000 File 1 reads. Enriching File 1 using 10 or more cycles of PCR increased the error rate between nts 28-32 and 167-176. Error rates for File 1 after DENSE, both in the elution and supernatant, remain largely unchanged.

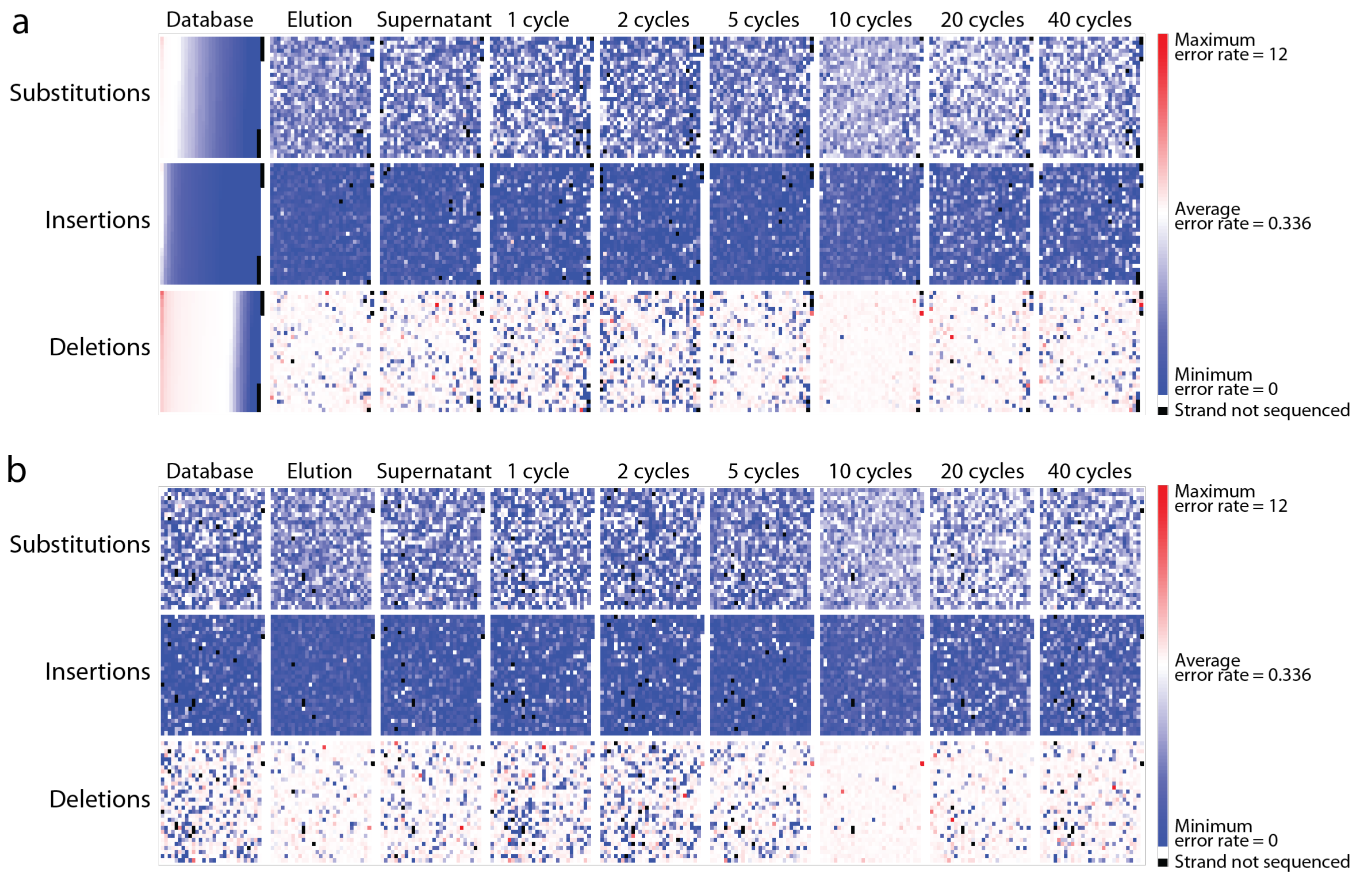


**Supplementary Figure 6. Error rate heatmaps of all File 1 strands after file access by different methods.** (left to right) Original database, biotin separation elution, biotin separation supernatant, 1 PCR cycle (without separation), 2 PCR cycles, 5 PCR cycles, 10 PCR cycles, 20 PCR cycles, and 40 PCR cycles. **(a)** In each row, strands are placed in order from highest to lowest rate of substitutions, insertions, or deletions, and the same order was maintained within each row. **(b)** To determine if particular strands have correlations in similar error types, all heat maps across all rows display the File 1 strands in the same order: Each unique strand is sorted by its index.

**Supplementary Table 1. Comparison of next generation sequencing and qPCR measurements of file ratios within samples indicate strong agreement between measurement methods.** All initial sample measurements were made using qPCR. Once NGS was conducted, results were compared to validate the accuracy of qPCR file ratio quantifications. Unknown sequences are those that did not fit the mapping criteria discussed in the Methods section.

| | File 1 | | File 2 | | File 3 | | File 4 | | File 5 | | unknown sequence |
|---|---|---|---|---|---|---|---|---|---|---|---|
| **Sample** | **NGS** | **qPCR** | **NGS** | **qPCR** | **NGS** | **qPCR** | **NGS** | **qPCR** | **NGS** | **qPCR** | **NGS** |
| Database | 12.23% | 11.00% | 16.12% | 19.00% | 13.96% | 17.00% | 26.59% | 33.00% | 24.99% | 26.00% | 6.10% |
| File 1 biotin elution (round 1) | 86.87% | 94.00% | 0.48% | 1.00% | 0.44% | 1.00% | 2.46% | 3.00% | 3.24% | 2.00% | 6.40% |
| File 1 biotin supernatant (round 1) | 18.10% | 32.00% | 10.30% | 16.00% | 9.55% | 19.00% | 26.99% | 18.00% | 32.34% | 15.00% | 2.70% |
| File 2 biotin elution | 0.25% | 1.00% | 94.17% | 95.00% | 0.12% | 0.00% | 1.20% | 2.00% | 1.60% | 2.00% | 2.60% |
| File 2 biotin supernatant | 4.14% | 6.00% | 27.27% | 29.00% | 6.67% | 11.00% | 26.59% | 24.00% | 33.03% | 29.00% | 2.30% |
| File 3 biotin elution | 0.60% | 1.00% | 1.20% | 2.00% | 86.15% | 92.00% | 2.87% | 3.00% | 3.81% | 3.00% | 5.30% |
| File 3 biotin supernatant | 4.99% | 5.00% | 9.18% | 13.00% | 28.91% | 41.00% | 24.61% | 22.00% | 30.53% | 20.00% | 1.80% |
| File 2 fluorescein eluiton | 0.25% | 0.00% | 92.29% | 85.00% | 0.49% | 0.00% | 1.24% | 8.00% | 1.64% | 7.00% | 4.00% |
| File 2 fluorescein supernatant | 6.38% | 6.00% | 26.54% | 10.00% | 11.83% | 8.00% | 24.64% | 34.00% | 28.11% | 43.00% | 2.50% |
| File 3 digoxigenin elution | 0.16% | 0.00% | 0.18% | 0.00% | 94.20% | 88.00% | 0.64% | 6.00% | 1.03% | 6.00% | 3.70% |
| File 3 digoxigenin supernatant | 4.92% | 5.00% | 8.48% | 5.00% | 34.71% | 9.00% | 21.95% | 38.00% | 25.58% | 43.00% | 4.30% |
| Repeated File 1 biotin elution (round 2) | 97.90% | 100.00% | 0.06% | 0.00% | 0.01% | 0.00% | 0.06% | 0.00% | 0.07% | 0.00% | 1.80% |
| Repeated File 1 biotin supernatant (round 2) | 48.30% | 56.00% | 1.70% | 2.00% | 1.70% | 4.00% | 20.20% | 18.00% | 26.30% | 20.00% | 1.80% |
| Repeated File 1 biotin elution (round 3) | 85.80% | 96.00% | 0.50% | 1.00% | 0.40% | 1.00% | 1.50% | 1.00% | 1.20% | 1.00% | 10.40% |
| Repeated File 1 biotin supernatant (round 3) | 59.80% | 64.00% | 0.40% | 1.00% | 0.40% | 1.00% | 16.30% | 19.00% | 19.10% | 15.00% | 4.10% |
| Repeated File 2 biotin elution (round 2) | 0.90% | 3.00% | 85.70% | 97.00% | 0.03% | 0.00% | 0.36% | 0.00% | 0.32% | 0.00% | 12.60% |
| Repeated File 2 biotin supernatant (round 2) | 8.70% | 12.00% | 25.50% | 31.00% | 2.20% | 4.00% | 27.10% | 27.00% | 33.60% | 26.00% | 2.80% |
| File 1 - 1 PCR cycle | 35.30% | 18.00% | 5.60% | 8.00% | 6.10% | 7.00% | 24.40% | 59.00% | 20.60% | 8.00% | 7.90% |
| File 1 - 2 PCR cycles | 50.20% | 39.00% | 4.10% | 8.00% | 4.80% | 11.00% | 18.60% | 36.00% | 16.30% | 6.00% | 6.00% |
| File 1 - 5 PCR cycles | 85.00% | 59.00% | 1.10% | 3.00% | 1.00% | 3.00% | 4.10% | 30.00% | 3.80% | 6.00% | 5.00% |
| File 1 - 10 PCR cycles | 95.50% | 29.00% | 0.50% | 6.00% | 0.80% | 7.00% | 0.70% | 22.00% | 0.90% | 36.00% | 1.70% |
| File 1 - 20 PCR cycles | 96.10% | 67.00% | 0.30% | 2.00% | 0.40% | 2.00% | 0.40% | 15.00% | 0.40% | 13.00% | 2.40% |
| File 1 - 40 PCR cycles | 94.70% | 95.00% | 0.40% | 0.00% | 0.30% | 0.00% | 0.40% | 0.00% | 0.40% | 4.00% | 3.90% |
